# Quantifying multivariate genotype-by-environment interactions, evolutionary potential and its context-dependence in natural populations of the water flea, *Daphnia magna*

**DOI:** 10.1101/2020.12.28.424558

**Authors:** Franziska S. Brunner, Alan Reynolds, Ian W. Wilson, Stephen Price, Steve Paterson, David Atkinson, Stewart J. Plaistow

## Abstract

Genotype-by-environment interactions (G x E) underpin the evolution of plastic responses in natural populations. Theory assumes that G x E interactions exist but empirical evidence from natural populations is equivocal and difficult to interpret because G x E interactions are normally univariate plastic responses to a single environmental gradient. We compared multivariate plastic responses of 43 *Daphnia magna* clones from the same population in a factorial experiment that crossed temperature and food environments. Multivariate plastic responses explained more than 30% of the total phenotypic variation in each environment. G x E interactions were detected in most environment combinations irrespective of the methodology used. However, the nature of G x E interactions was context-dependent and led to environment-specific differences in additive genetic variation (G-matrices). Clones that deviated from the population average plastic response were not the same in each environmental context and there was no difference in whether clones varied in the nature (phenotypic integration) or magnitude of their plastic response in different environments. Plastic responses to food were aligned with additive genetic variation (*g*max) at both temperatures, whereas plastic responses to temperature were not aligned with additive genetic variation (*g*max) in either food environment. These results suggest that fundamental differences may exist in the potential for our population to evolve novel responses to food versus temperature changes, and challenges past interpretations of thermal adaptation based on univariate studies.

## INTRODUCTION

A genotype-by-environment interaction (G x E) occurs whenever genotypes differ in the way that their trait values change across environments (Saltz *et al*., 2018), or in other words, when there is genetic variation in phenotypic plasticity (Gillespie and Turelli, 1989; Schlichting and Pigliucci, 1993; Pigliucci and Preston, 2004). G x E interactions are critical for understanding population responses to environmental change because they alter the expression of heritable phenotypic variation between different environments (Hoffmann and Merila, 1999; Gibson and Dworkin, 2004; Schlichting, 2008; Ledón-Rettig *et al*., 2014; Wood and Brodie, 2016). Moreover, they underpin the evolution of phenotypic plasticity in natural populations (Pigliucci, 2005; Levis and Pfennig, 2016; Oostra *et al*., 2018; Fox *et al*., 2019). When environments change rapidly, plasticity may be the only response possible. Consequently, G x E interactions and the potential to evolve adaptive plastic responses are crucial for population survival in the face of climate change (Chevin *et al*., 2010; Snell-Rood *et al*., 2018).

There is increasing evidence that much of the phenotypic change associated with climate change in wild populations is attributable to phenotypic plasticity (Chevin and Hoffmann, 2017). Phenotypic variation explained by G x E interactions is expected to be larger in novel or extreme environments because plastic responses will not yet have been exposed to selection, leading to the release of what is termed cryptic genetic variation (Gibson and Dworkin, 2004; Schlichting, 2008; Mcguigan and Sgrò, 2009; Ledón-Rettig *et al*., 2014; Paaby and Rockman, 2014). But in environments that have always been part of a population’s evolutionary history, long-term selection for an optimal plastic response may have depleted G x E interactions. For example, in the African savannah butterfly, *Bicyclus anynana*, genetic variation for seasonal plasticity was almost absent despite considerable additive genetic variation for trait means (Oostra *et al*., 2018). When G x E interactions are absent, traits can evolve but trait plasticity cannot, increasing a population’s susceptibility to changes in the frequency of extreme events such as heatwaves, droughts and floods (Easterling *et al*., 2000), or changes in the reliability of existing environmental cues (Reed *et al*., 2010; Oostra *et al*., 2018; Bonamour *et al*., 2019).

Theory assumes that G x E interactions exist in natural populations (Via and Lande, 1985; Lande, 2009), and G x E interactions are frequently demonstrated in plant studies (Des Marais *et al*., 2013) and laboratory studies (Vieira *et al*., 2000; Valdar *et al*., 2006; Ingleby *et al*., 2010; Plaistow and Collin, 2014). But evidence for G x E interactions in wild animal populations is sparse (Nussey *et al*., 2005a, b). Moreover, other recent studies haven’t detected G x E interactions (Brommer *et al*., 2008; Charmantier and Gienapp, 2014; Hayward *et al*., 2018; Oostra *et al*., 2018). G x E interactions are normally assessed in a single trait (Hayward *et al*., 2018) but plastic responses typically involve coordinated shifts in many traits at the same time (Plaistow and Collin, 2014), and univariate studies do not distinguish differences in the magnitude of a plastic response from differences in the nature of a plastic response (Chun *et al*., 2007; Collyer and Adams, 2007; Adams and Collyer, 2009; Plaistow and Collin, 2014). Figure 1 shows G x E interactions in two hypothetical populations that have the same average multivariate plastic response to an environmental contrast. Differences in the magnitude of multivariate plastic responses will create G x E interactions that generate most additive genetic variation (gmax) in alignment with the average plastic response (see Fig. 1A). But differences in the nature of a multivariate plastic response generate additive genetic variation (gmax) that is not aligned with the population average plastic response (see Fig 1B). Since populations are expected to have increased additive genetic variation along the axis of the average plastic response (Lande, 2009; Draghi and Whitlock, 2012; Noble *et al*., 2019) the type of multivariate G x E interactions in a population may have important consequences for a population’s ability to rapidly adapt to an environmental change.

**Figure 1.**
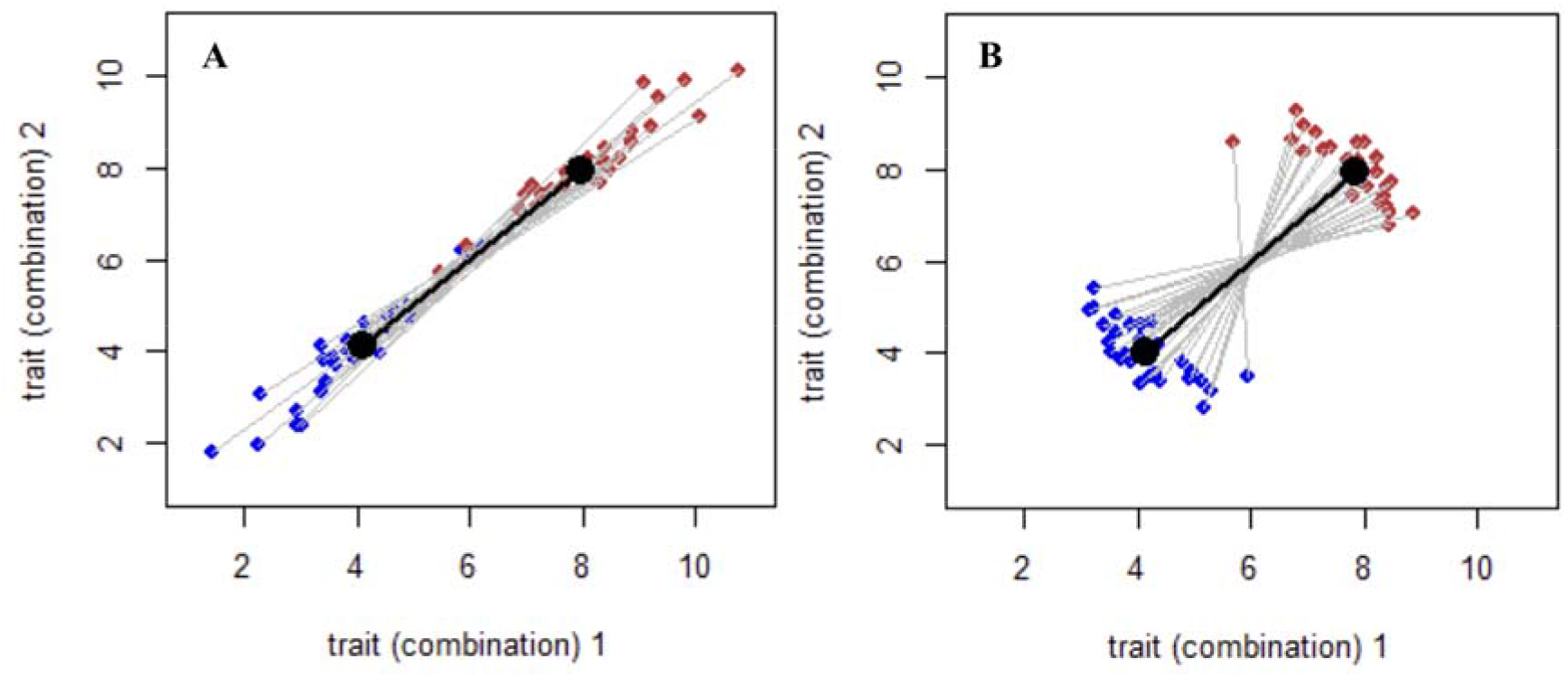
G x E interactions in two hypothetical populations that have the same average plastic response. Differences in the magnitude of plastic responses (A) generate additive genetic variation that is aligned with the average plastic response whereas differences in the nature of plastic responses (B) generate additive genetic variation that is not aligned with the average plastic response.

Multivariate G x E interactions can be tested for using a character-state approach (Via and Lande, 1985) that uses Bayesian Markov chain Monte Carlo (MCMC) mixed models to compare the volume, shape and orientation of G-matrices estimated for the same genotypes reared in two or more different environments. The volume of the G-matrix characterises the amount of clonal genetic variation (V_G_) that selection can act on, whereas the shape defines whether the variation is attributable to few or many traits, and the orientation, defined by *g*max, identifies the traits associated with the most clonal genetic variation (V_G_) (Robinson and Beckerman, 2013; Lind *et al*., 2015; Reger *et al*., 2017). Alternatively, multivariate G x E interactions can be detected by testing for differences in the length or angle of reaction norms generated by measuring a multivariate phenotype of genotypes or families in two or more environments (Collyer and Adams, 2007; Plaistow and Collin, 2014). Irrespective of the approach used, multivariate G x E interactions are normally only quantified in response to a single environmental variable despite the fact that environments are often complex and have multiple dimensions that vary simultaneously (Westneat *et al*., 2019). As a result, we do not yet know if multivariate G x E interactions are context-dependent, or if the evolved plastic response to one environmental variable has implications for a population’s ability to evolve a response to a different environmental variable.

Clonal organisms such as *Daphnia* are ideal systems for studying G x E interactions because it is easy to separate genetic and non-genetic influences, and large numbers of genetically identical individuals can be reared simultaneously across different environments in parental and offspring generations (Harris *et al*., 2012; Miner *et al*., 2012). In order to better understand the evolutionary significance of multivariate G x E interactions in natural populations we isolated 43 genotypes (clones) of *Daphnia magna* from a single population and measured their multivariate plastic response to temperature in different food environments and their multivariate plastic response to food at different temperatures. We used a character-state approach and the tools developed in (Robinson and Beckerman, 2013) to test for genetic variation in multivariate plastic responses and its context-dependence and to compare the alignment of average plastic responses with *g*max (Lind *et al*., 2015; Noble *et al*., 2019; Radersma *et al*., 2020). We then used a reaction norm approach to determine the source of multivariate G x E interactions in each environmental context (differences in magnitude or phenotypic integration) and the number and identity of clones that diverged from the average multivariate plastic response in each case (Collyer and Adams, 2007). We compared our results to two reference populations from a higher and a lower latitude in order to assess the generality of our findings.

## METHODS

### (a) Experimental animals

*Daphnia magna* clones from the UK were collected as resting eggs from Brown Moss, a shallow wetland in Shropshire (52°57’01”N 2°39’05”W, National Grid Reference SJ 562395) in July 2016. The eggs were stored in total darkness for a 3-month period at 4°C before being hatched out at 21°C on a 14:10 light: dark cycle. The 5 French clones were collected in 2007 from small pools in the Camargue in the South of France (van Doorslaer *et al*., 2009a) and the 8 Danish clones were collected from Lake Ring in 2000 (Michels, 2007). The clones were maintained as laboratory stock cultures in a controlled temperature incubator at 21°C ± 1°C on a 14:10 light: dark cycle. Animals were kept in 200ml glass jars containing 150 ml of artificial pond water media (ASTM) enriched with additional organic extracts (AQ Xtract 30, Wilfrid Smith, UK) (Baird *et al*., 1989). The jars were fed high food three times a week (200,000 cells ml^-1^ of batch-cultured *Chlorella vulgaris*, quantified with a haemocytometer) and changed to fresh media once a fortnight.

### (b) Experimental set-up

Prior to starting the experiment, three females from each clone were isolated from stocks and kept individually on *ad libitum* food (2×10^5^ cells ml^-1^ day^-1^ of batch-cultured *C. vulgaris*) until they produced a clutch. From that clutch, one offspring was randomly selected and reared individually in separate 200 ml jars fed for 2 generations to reduce possible maternal effects (Plaistow and Collin, 2014). In the 3rd generation, 3 - 8 second clutch neonates per clone were randomly allocated to 4 different experimental rearing treatments generated by crossing food levels (2×10^5^ and 4×10^4^ *C. vulgaris* cells ml^-1^ day^-1^) and temperature (24°C and 18°C). The jars were observed daily and transferred to a jar with fresh food and media every other day until they themselves dropped their second clutch. Experiments were performed in 5 temporal blocks between January 2017 and March 2018, each clone being assayed in multiple blocks.

### (c) Life history traits

All 856 individuals in the study were photographed as neonates, upon reaching maturity (first eggs in the brood pouch) and upon releasing their second clutch, using a Canon EOS 350D digital camera connected to a Leica MZ6 dissecting microscope. The number of neonates each individual produced in their 1^st^ and 2^nd^ clutch were counted and 5 neonates in each clutch were photographed to obtain an estimate of offspring size. In all cases, body size was measured as the distance from the top of the head to the base of the tail spine using the image analysis software, ImageJ version 1.45s (Rasband, 1997). After mothers released their second clutch, their thermal tolerance (CT_max_) was assayed as described by (Geerts *et al*., 2015), allowing animals an acclimation period of 15 mins at 21°C followed by a ramping period of 40s/°C from 21°C to 50°C. So in total we measured thermal tolerance and 8 life-history traits for each individual: length at maturity, length at second clutch, age at maturity, age at second clutch, juvenile growth rate ((length at maturity - length as neonate)/ age at maturity), adult growth rate ((length at second clutch - length at maturity)/(age at second clutch-age at maturity)), average fecundity (across clutches 1 and 2), and average offspring size (across clutches 1 and 2).

### (d) Statistical analyses

To test for multivariate GxE interactions and their context-dependence we estimated environment specific variance-covariance matrices (G) using a Bayesian MCMC multivariate mixed model (Hadfield, 2010). We then used the tools developed in (Robinson and Beckerman, 2013) to compare the volume, shape and orientation of the G-matrices generated by the two temperature treatments when experiencing high food and low food, and the G-matrices generated by the food treatments when experiencing high and low temperatures. All trait variables were centred and scaled to s.d. = 1. We used parameter-expanded priors, and models were fitted with a burn-in of 50,000 and sampling that produced 1000 estimates of the joint posterior distribution from more than 500,000 iterations of the chains. All models were checked for autocorrelation in the chains. To determine the source of multivariate G x E interactions in each environmental context (differences in magnitude or phenotypic integration) we analysed the 8 life history traits and the thermal tolerance of each individual using perMANOVAs with temperature, food level and clone and all their interactions fitted as fixed factors and temporal block fitted as a random factor. We used marginal R^2^ values from the models to determine the proportion of phenotypic variance attributable to different model components. The multivariate plastic response of each clone to changes in food and temperature were quantified as phenotypic change vectors of scaled phenotypic data (9 traits: 8 life history variables and thermal tolerance CT_max_) following (Collyer and Adams, 2007; Plaistow and Collin, 2014). Separate vectors were fitted for plastic responses to food and temperature in each environmental context, i.e. plasticity in response to temperature was quantified once in the low food environment and once in the high food environment and *vice versa*. The magnitude of the plastic response for each clone was calculated as the Euclidian length of the phenotypic change vector while the nature of the plastic response was calculated as the angle between a specific clone’s plastic response and the mean response within the population (Collyer and Adams, 2007; Plaistow and Collin, 2014). The magnitude and nature of each clone’s multivariate plastic response were then compared to the population mean response using a permutation procedure, where the actual deviation from the average response was compared to deviations generated by random vector pairs iteratively sampled from the data 9999 times (Collyer and Adams, 2007). The significance levels for these tests were adjusted by the Benjamini-Yekutieli method to α=0.024 to correct for testing each clone in 4 separate tests (magnitude and phenotypic integration each in 2 environments). Differences in multivariate phenotypes and reaction norms were visualised by projecting multivariate trait means onto the first two principal component axes of a PCA using all 8 life-history traits and thermal tolerance (Chun *et al*., 2007; Plaistow and Collin, 2014). The same analyses described above were then also used to compare the reaction norm variation in the UK population with the reaction norms of 5 clones collected from a single population from the south of France (lower latitude) and 8 clones collected from a single population in Denmark (higher latitude). All of the analyses were conducted in R version 3.5.2 (Team, 2019) using the ‘vegan’ package (Oksanen, J. *et al*., 2018).

## RESULTS

### (a) A comparison of G-matrices in different environmental contexts

Temperature had no effect on the volume or shape of the variance-covariance matrices (G) in either a high food or a low food environment (Table 1, Fig. 2 A,B). But temperature altered the orientation of the variance-covariance matrix (G) in both food environments. In the high food environment, additive genetic variation in clutch size explained *g*max at 24°C but clones with larger clutches also developed at a faster rate at 18°C. Similarly, in the low food environment additive genetic variation in CTmax explained *g*max at 24°C but clones with a lower CTmax matured faster at 18°C (Table 1, Fig. 2 A,B).

**Table 1.** Matrix comparison statistics for

| Plasticity (context) | Metric | Mode | Lower 95% CI | Upper 95% CI | Probability |
| --- | --- | --- | --- | --- | --- |
| **Temperature (high food)** | Var gmax Diff | -0.040 | -0.292 | 0.247 | NA |
|  | **Angle between *gmax*** | **39.683** | **22.104** | **69.414** | **<0.05** |
|  | Prob-Vol diff | 0.000 | 0.000 | 0.000 | NA |
|  | Sum-Vol diff | 0.711 | -17.000 | 9.602 | NA |
| **Temperature (Low food)** | Var gmax Diff | -0.070 | -0.327 | 0.203 | NA |
|  | **Angle between *gmax*** | **41.122** | **23.731** | **63.229** | **<0.05** |
|  | Prob-Vol diff | 0.000 | 0.000 | 0.000 | NA |
|  | Sum-Vol diff | 0.107 | -6.499 | 6.803 | NA |
| **Food (24°C)** | Var gmax Diff | -0.059 | -0.283 | 0.233 | NA |
|  | Angle between *gmax* | 42.746 | 18.054 | 75.971 | 0.158 |
|  | Prob-Vol diff | 0.000 | 0.000 | 0.000 | NA |
|  | Sum-Vol diff | 5.950 | -0.403 | 18.220 | NA |
| **Food (18°C)** | Var gmax Diff | -0.090 | -0.300 | 0.221 | NA |
|  | **Angle between *gmax*** | **56.184** | **36.123** | **78.765** | **<0.05** |
|  | Prob-Vol diff | 0.000 | 0.000 | 0.000 | NA |
|  | **Sum-Vol diff** | **8.730** | **0.068** | **21.674** | **<0.05** |

**Figure 2.**
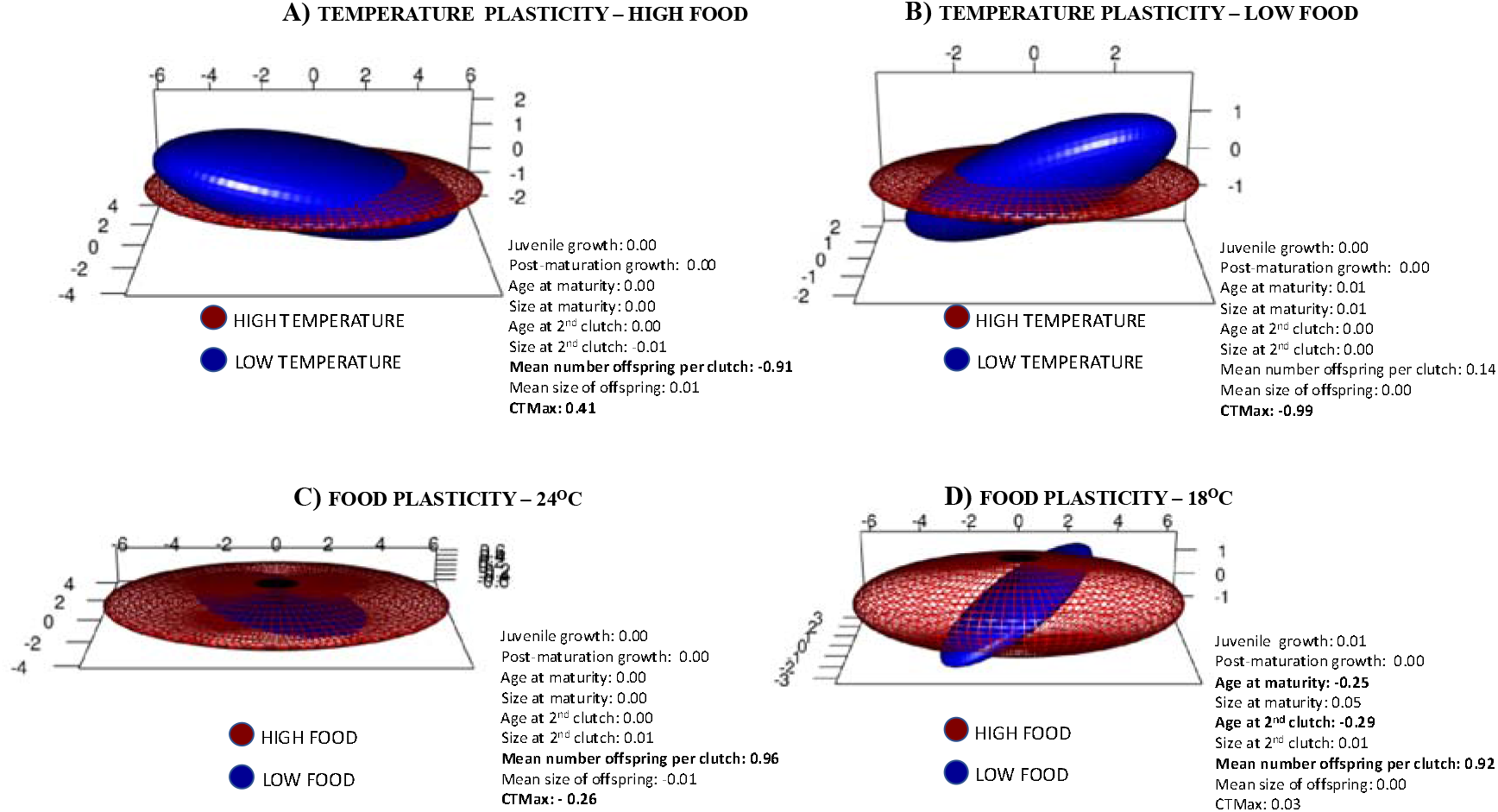
Genetic variance–covariance matrix visualizations for each treatment within each environmental context. The volume of the three-dimensional hull represents the amount of additive genetic variance whereas the shape and rotation reflect changes in covariance between traits. The loadings for PC1 represent the contributions of traits towards *g*max in each environmental context.

Food had no effect on the volume, shape or orientation of the variance-covariance matrices (G) at 24°C (Table 1, Fig. 2 C). But at 18°C, the volume of the high food G-matrix was significantly larger than the volume of the low food G-matrix (Table 1, Fig. 2C) demonstrating that there is more additive genetic variation in traits such as mean clutch size and development time in high food environments. Moreover, the orientation of the low food matrix was significantly rotated towards differences in age at maturity such that clones with fewer offspring per clutch also tended to have higher CTmax and slower development rates (Table 1, Fig. 2D). In order to explore the cause of the multivariate G x E’s in more detail we compared clonal differences in the nature and magnitude of plastic responses to temperature and food.

### (b) Multivariate plastic responses to temperature in different food environments

The multivariate plastic response to temperature explained just under 40% of the phenotypic variation in both the high food environment (36.9%) and the low food environment (35.8%) and was characterised by increased juvenile growth rates, earlier maturation and earlier production of the second clutch at higher temperatures (Table 2; Fig. 3A-C). Additive clonal variation explained slightly more life-history trait variation in the high food environment (23.2%) compared to the low food environment (18.5%). A significant G x E interaction explained 10.8% of phenotypic variation in the high food environment and 9.8% in the low food environment (Table 2; Fig. 3 B,C). Across both food environments, permutation tests revealed that 18 out of 43 clones (41.8%) deviated significantly from the population average multivariate response to temperature in one way or another. In the high food environment 5 clones deviated in the magnitude of their plastic response, 4 clones deviated in the nature of their plastic response and 3 clones deviated in both the magnitude and the nature of their plastic response. In the low food environment, 2 clones deviated in the magnitude of their plastic response, 5 clones deviated in the nature of their plastic response and 2 clones deviated in both the magnitude and the nature of their plastic response. Only 3 of the 18 clones deviated from the population average plastic response in some way in both environments (see Fig. 3D) highlighting the context-dependence of multivariate G x E interactions in response to temperature. Permutation test outcomes for each clone’s plasticity values against population means can be found in the supplementary data.

**Table 2.**
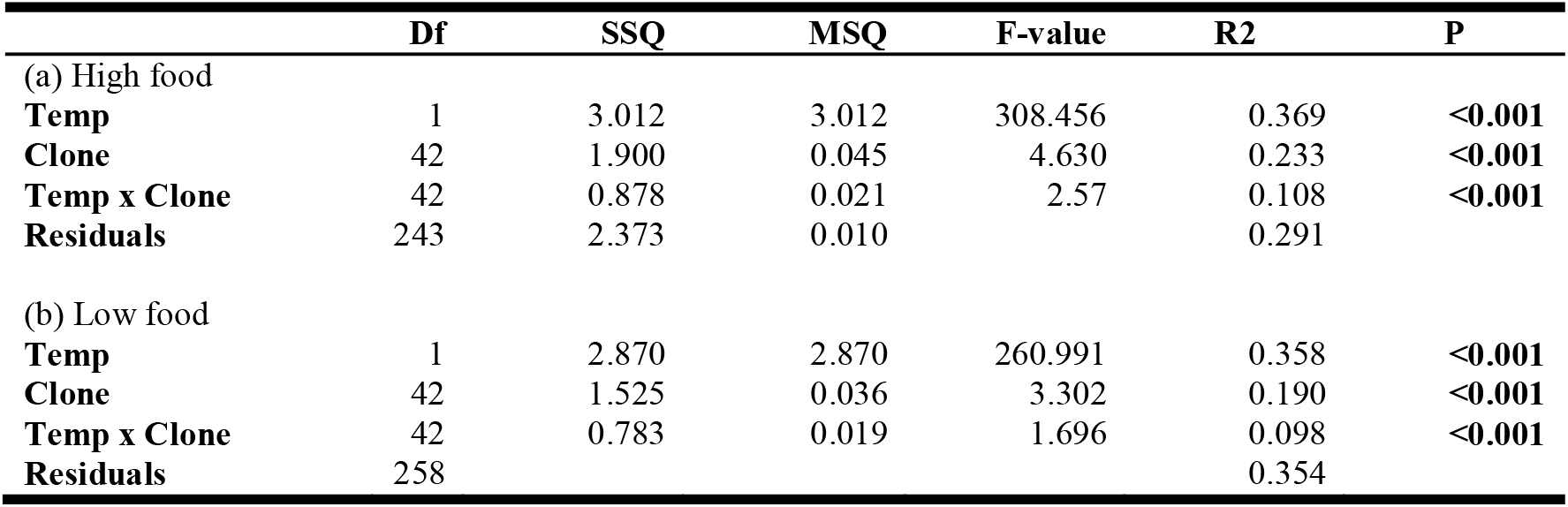
Multivariate plastic responses to temperature in high and low food environments.

**Figure 3:**
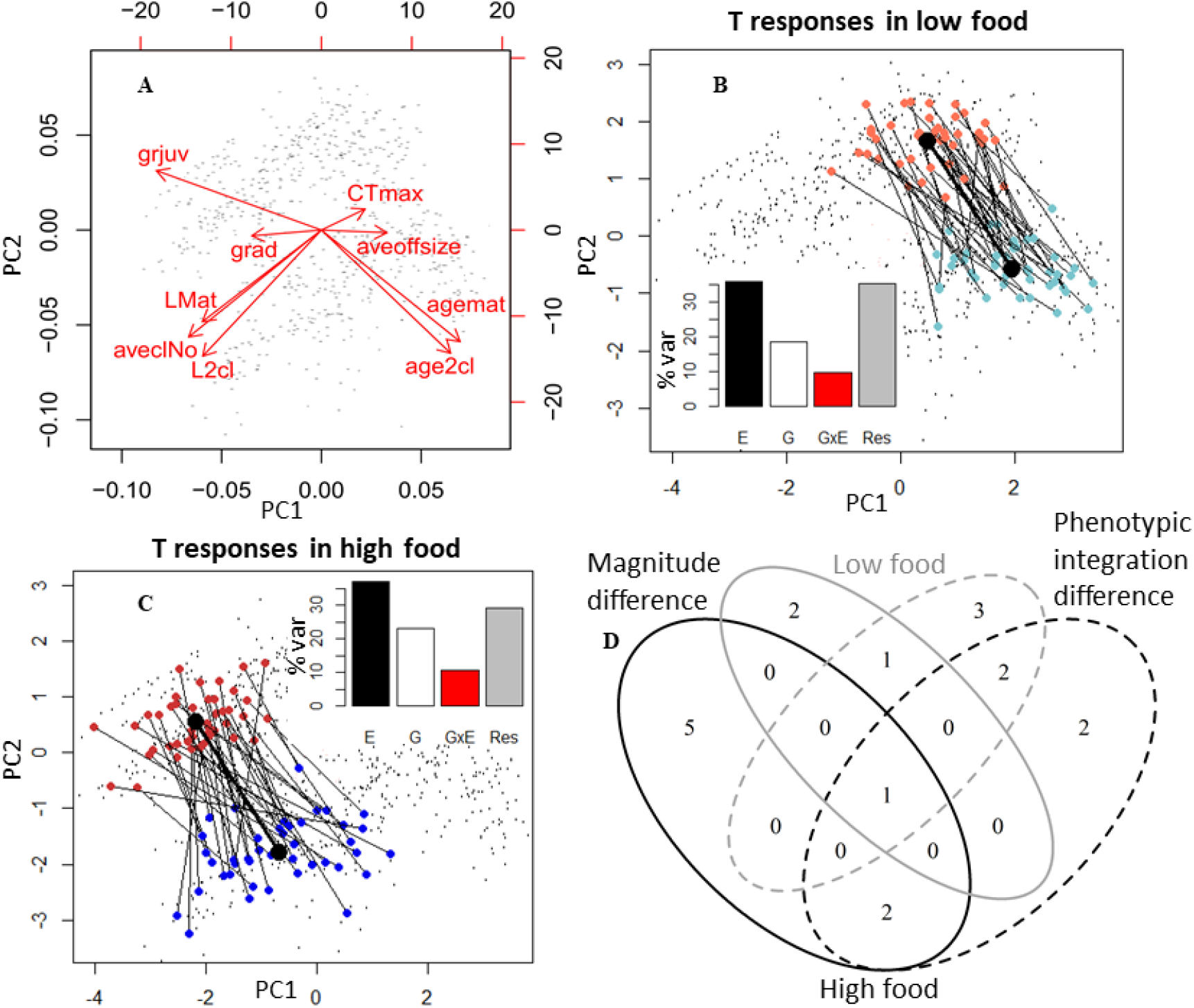
Variable plastic responses to temperature within the UK population. (A) The first two Principal Component Axes summarize the multivariate trait space as shown by PCA biplot. agemat = age at maturity, age2cl = age at second clutch, LMat = length at maturity, L2cl = length at second clutch, aveclNo = average fecundity across clutches 1 and 2, aveoffsize = average offspring size, grjuv = juvenile growth rate, grad = adult growth rate, CTmax = temperature tolerance. Reaction norms in response to temperature for each clone are shown in the low food context (B) and in the high food context (C) within the same multivariate space. Average clonal phenotypes in each environment are indicated in dark red/light red for 24°C and dark blue/light blue for 18°C. Insets show components of variation estimated from marginal R^2^ in perMANOVA models. E= (temperature) environment, G = genotype, GxE=plasticity variation, Res= Residuals. (D) Differences from population means for each clone are summarised in a Venn diagram to show the overlap in clones for different outlier tests. Black and grey outlines indicate high food and low food environment, respectively; solid and dashed lines indicate differences from population mean response in magnitude or phenotypic integration, respectively.

### (c) Multivariate plastic responses to food in different temperatures

The multivariate plastic response to food explained 40.1 % of the phenotypic variation at 24°C but only 33.8% of the phenotypic variation at 18°C. Individuals reared in high food environments matured at larger sizes, were larger at second clutch and had more offspring per clutch compared to individuals reared on low food at both temperatures (Table 3, Fig. 4A-C) but additive clonal variation explained more phenotypic variation at 18°C (26.9%) than at 24°C (19.6%). Similarly, a significant G x E interaction (Table 2; Fig. 4 B,C) also explained more phenotypic variation at 18°C (9.6%) than at 24°C (6.7%). In total, 16 of the 43 clones (37.2%) had a multivariate plastic response to food that deviated significantly from the population average in one way or another (see Fig. 4D). At 18°C, 5 clones deviated in the magnitude of their plastic response, 5 clones deviated in the nature of their plastic response and 2 clones deviated in both the magnitude and the nature of their plastic response. In comparison, at 24°C only 2 clones each deviated in the magnitude and in the nature of their plastic response and no clones deviated in both the magnitude and the nature of their plastic response. None of the 16 clones deviated from the population average plastic response both at 18°C and 24°C (see Fig. 4D), highlighting the context-dependence of multivariate G x E interactions in response to food. Permutation test outcomes for each clone’s plasticity values against population means can be found in the supplementary data.

**Table 3.** Multivariate plastic response to food at high and low temperatures.

|  | **Df** | **SSQ** | **MSQ** | **F-value** | **R2** | **P** |
| --- | --- | --- | --- | --- | --- | --- |
| (a) High temperature |  |  |  |  |  |  |
| **Food** | 1 | 3.081 | 3.081 | 303.041 | 0.401 | **<0.001** |
| **Clone** | 42 | 1.499 | 0.036 | 3.510 | 0.195 | **<0.001** |
| **Food x Clone** | 42 | 0.514 | 0.012 | 1.203 | 0.067 | **<0.001** |
| **Residuals** | 255 | 2.592 | 0.010 |  | 0.337 |  |
| (b) Low temperature |  |  |  |  |  |  |
| **Food** | 1 | 3.428 | 3.428 | 297.379 | 0.338 | **<0.001** |
| **Clone** | 42 | 2.907 | 0.069 | 6.005 | 0.287 | **<0.001** |
| **Food x Clone** | 42 | 0.969 | 0.023 | 2.001 | 0.096 | **<0.001** |
| **Residuals** | 246 |  |  |  | 0.280 |  |

**Figure 4.**
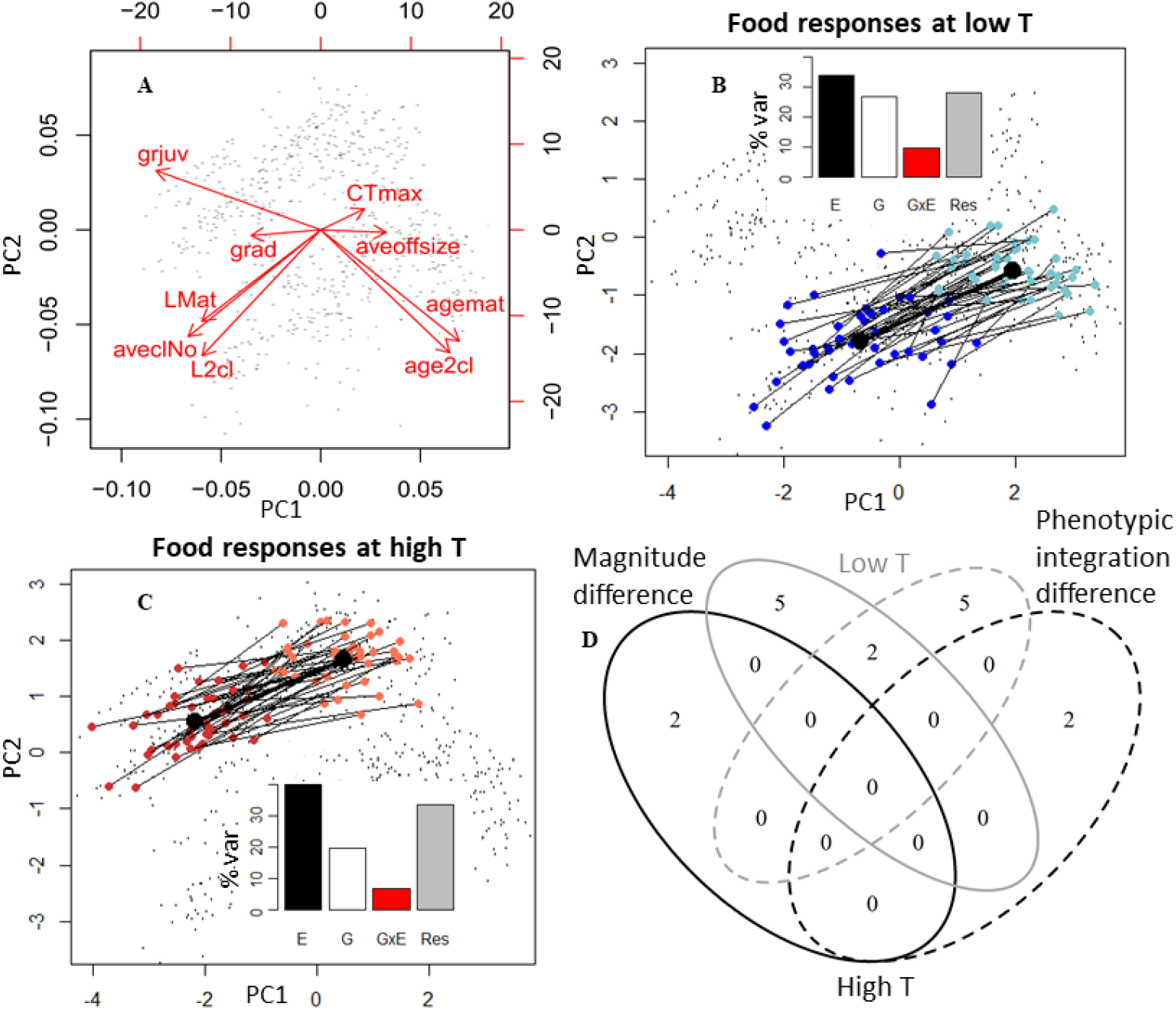
Variable plastic responses to resources within the UK population. (A) The first two Principal Component Axes summarize the multivariate trait space as shown by PCA biplot. agemat = age at maturity, age2cl = age at second clutch, LMat = length at maturity, L2cl = length at second clutch, aveclNo = average fecundity across clutches 1 and 2, aveoffsize = average offspring size, grjuv = juvenile growth rate, grad = adult growth rate, CTmax = temperature tolerance. Reaction norms in response to resources for each clone are shown in 18°C context (B) and in the 24°C context (C) within the same multivariate space. Average clonal phenotypes in each environment are indicated in dark red/dark blue for high food and light red/light blue for low food. Insets show components of variation estimated from marginal R^2^ in perMANOVA models. E= (temperature) environment, G = genotype, GxE=plasticity variation, Res= Residuals. (D) Differences from population means for each clone are summarised in a Venn diagram to show the overlap in clones for different outlier tests. Black and grey outlines indicate high food and low food environment, respectively; solid and dashed lines indicate differences from population mean response in magnitude or phenotypic integration, respectively.

### (d) Multivariate plasticity in different populations

When we compared our UK population with a smaller subset of clones from a lower latitude population (South of France) and a higher latitude population (Denmark), we found no evidence that the average plastic response to temperature differed between populations despite their different latitudinal origins (Pop x Temp interaction, F_1,844_=-0.0017, P=1, FigS1, Table S1). When exposed to a higher temperature, individuals in all three populations grew faster and matured earlier (see Fig. 5). However, the average plastic response to food levels did differ between populations (Pop x Food interaction, F_2,844_=0.0025, P=0.016). In high food, individuals in all populations matured at larger sizes and produced larger clutches of offspring but less so in the French population (see Fig. 5K). The proportion of phenotypic variation explained by different model components was analysed for each population separately. Average multivariate plastic responses to food and temperature explained between 39.5 − 57.6% of the phenotypic variation in the French and Danish populations in different contexts. Clone-specific multivariate plastic responses to food and temperature were both context-dependent in the French population (see Table S2b) as in the UK population (see above). G x E interactions in response to temperature explained 12.4% of the total phenotypic variation in high food but only 6.5% in low food. Whereas G x E in response to food explained 13.2 % of the phenotypic variation in low temperature but only 6.5% of the phenotypic variation in the high temperature. In the Denmark population, plastic responses to temperature were indistinguishable between clones (see Table S2c?), whereas a significant G x E in response to food (see Table S2c) explained 8.4% of the phenotypic variation in low temperatures and 9.7% of the phenotypic variation in high temperatures.

**Figure 5.**
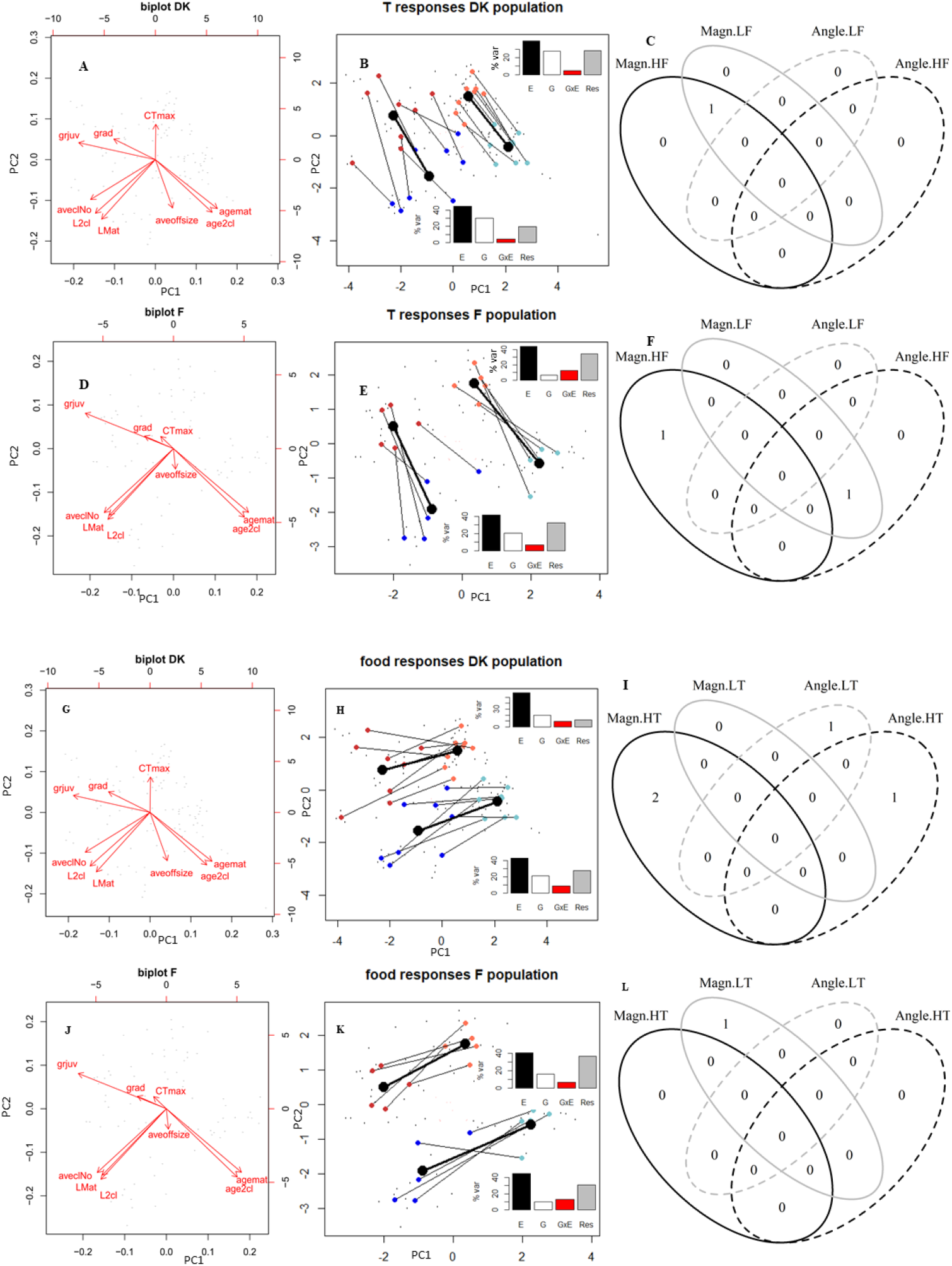
Plastic responses to temperature and resources within two reference populations. (A, D, G, J) The first two Principal Component Axes summarize the multivariate trait space as shown by PCA biplot. agemat = age at maturity, age2cl = age at second clutch, LMat = length at maturity, L2cl = length at second clutch, aveclNo = average fecundity across clutches 1 and 2, aveoffsize = average offspring size, grjuv = juvenile growth rate, grad = adult growth rate, CTmax = temperature tolerance. Reaction norms in response to temperature for each clone in the Danish (B) and French (E) populations are shown in high food context (dark red/ dark blue) in low food context (light red/light blue) within the same multivariate plot. Reaction norms in response to food for each clone in the Danish (H) and French (K) populations are shown in high temperature context (dark red/ light red) and in low temperature context (dark blue/light blue) within the same plot. Population average plastic responses are shown by solid black lines. Insets show components of variation estimated from marginal R^2^ in perMANOVA models. E= (temperature) environment, G = genotype, GxE=plasticity variation, Res= Residuals. (C, F, I, L) Differences from population means for each clone are summarised in a Venn diagram to show the overlap in clones for different outlier tests.

## DISCUSSION

Theory assumes that G x E interactions exist in natural populations (Via and Lande, 1985; Lande, 2009; Chevin and Hoffmann, 2017) but empirical evidence is equivocal (Hayward *et al*., 2018) and not always easy to interpret when G x E interactions are measured as univariate responses to a single environmental gradient. We compared the multivariate plastic responses to food and temperature environments for 43 *Daphnia magna* clones collected from the same population. Average multivariate plastic responses to food-temperature environments explained three times more phenotypic variation than genetic variation in traits (G), or genetic variation in plasticity (G x E), supporting the idea that the phenotypic change attributable to plasticity in wild populations is higher than previously thought (Charmantier *et al*., 2008; Gienapp *et al*., 2008; Merilä and Hendry, 2014). Consequently, it is imperative that we understand how plasticity evolves and influences a population’s ability to cope with environmental change (Chevin *et al*., 2010). G x E interactions are critical as plasticity cannot evolve in their absence (Pigliucci, 2005; Hayward *et al*., 2018; Oostra *et al*., 2018; Fox *et al*., 2019).

Using a reaction norm approach we detected multivariate G x E interactions in all three of the populations examined. Moreover, in our focal population we detected multivariate G x E interactions in almost all of the environmental contexts we examined irrespective of whether we used a character-state approach (Via and Lande, 1985; Robinson and Beckerman, 2013), or a reaction norm approach (Collyer and Adams, 2007). But it remains to be seen whether the 10% or less of phenotypic variation explained by G x E in a laboratory environment is sufficient to fuel the evolution of plastic responses in a natural environment. Recent studies have concluded that G x E interactions are not detectable in wild populations (Brommer *et al*., 2008; Charmantier and Gienapp, 2014; Hayward *et al*., 2018; Oostra *et al*., 2018) but these studies only tested for univariate G x E interactions arising from differences in the magnitude of a plastic response. Our multivariate study also allowed us to test for differences in the nature of a plastic response, another potentially important source of G x E interaction (Plaistow and Collin, 2014). Food had no effect on the volume, shape or orientation of variance-covariance matrices (*G*) at high temperatures, suggesting that there was no genetic variation in the way that different clones responded to food at high temperatures. This is further supported by the reduced phenotypic variance explained by food plasticity G x E interactions at 24°C compared to 18°C, and fewer clones deviating from an average plastic response (Fig. 4). But in all other environmental contexts differences in the orientation of variance-covariance matrices (G) in different environments was observed, meaning that the combination of traits that expressed the most additive variation changed. For example, for temperature plasticity in a high food environment, most of the additive genetic variation (*g*max) was attributable to a trade-off between the mean number of offspring in each clutch and CT max at 24°C, but at 18°C the trade-off also involved differences in age at maturity (see Fig. 2A,B). For food plasticity at 18°C, the orientation of the variance-covariance matrices (G) was significantly different, but there was also a significant reduction in the volume of variance-covariance matrix in the low food environment, meaning that the population’s evolutionary potential was reduced in low food compared to high food.

We hypothesised that context-dependent multivariate G x E interactions might be explained by differences in the number and identity of clones whose multivariate reaction norms diverged in either magnitude or phenotypic integration from the average multivariate plastic response. But there was no pattern in the number of clones that differed in the magnitude or nature of their multivariate plastic response in different environments. Clones that deviated from the population average plastic response were not the same in each environment, suggesting that the outcome of selection in different environments must also be context-dependent, which may help explain why clonal variation is often maintained in natural populations (Hebert and Crease, 1980; Weeks and Hoffmann, 2008).

Studies normally only consider the evolution of plastic responses to a single environmental variable (Dennis *et al*., 2011; Plaistow and Collin, 2014; Westneat *et al*., 2019). The context-dependence of the G x E’s observed in this study, and the effect that context-dependence had on evolutionary potential, suggests that it may be important to consider the evolution of plastic responses to more complex environmental cues if we want to understand how plasticity contributes to the demography and extinction risk of populations (Hoffmann and Sgrò, 2011; Merilä and Hendry, 2014; Chevin and Hoffmann, 2017). Context-dependent G x E interactions could be explained by plastic responses to one environmental variable altering plastic responses to another environmental variable. For example, in our study multivariate G x E interactions at 18°C but not at 24°C could arise because development is faster at higher temperatures and constrains the effect that food plasticity has on traits such as body size and clutch size (Atkinson, 1994; Geerts *et al*., 2015) Similarly, a reduced evolutionary potential in low food at 18°C could be because low food constrains the expression of life-history traits, making G x E interactions less detectable. However, context-dependent evolutionary potential can also arise because the strength or nature of selection varies between environments. Selection-by-environment interactions are common and are widely reported (Wood and Brodie, 2016; Hayward *et al*., 2018). Alternatively, some environments may be more common than others over a population’s evolutionary history, allowing more occasions for selection to optimise multivariate responses to that particular environment (Chevin and Hoffmann, 2017).

Quantifying G x E interactions is important for understanding the potential for plasticity to evolve in populations. But it is how the detected G x E influences heritable trait variation in each environment that will ultimately determine how the genotype – phenotype relationship interacts with selection (Draghi and Whitlock, 2012). The adaptive plastic responses that organisms have evolved can be viewed as a kind of developmental bias that converts environmental and genetic cues into variation in the plastic response (Draghi and Whitlock, 2012). As a result, the traits that contribute to an evolved plastic response are predicted to be the same traits that show the most additive genetic variation (*g*max) in populations introduced to new environments (Draghi and Whitlock, 2012; Noble *et al*., 2019; Radersma *et al*., 2020). Interestingly, we found a close relationship between the average plastic response to food and *g*max at 18°C and 24°C (Fig. 4) but no relationship between the average plastic response to temperature and *g*max in either food environment (Fig. 3). This difference could arise if temperature has exerted a stronger selection pressure than food over evolutionary time and was therefore more effective at removing additive genetic variation. Alternatively, food environments may hide genetic variation from selection. Harsh environments are often assumed to exert the strongest selection pressures on a population (Hayward *et al*., 2018), but harsh environments that reduce evolutionary potential, or have little demographic consequence, can also shield additive genetic variation from selection (Jong and Behera, 2002; Chevin and Hoffmann, 2017). In *Daphnia*, low food environments contribute little to demographic change (Heugens *et al*., 2006) and reduce evolutionary potential (this study). This may explain why evolutionary responses to temperature manipulations were more likely in populations that were not food limited (van Doorslaer *et al*., 2009b; De Meester *et al*., 2011).

Irrespective of the causes of context-dependent differences in evolutionary potential, our results demonstrate that *Daphnia magna* populations have maintained genetic potential to evolve adaptive responses to resources compared to their capacity to evolve adaptive responses to temperature. This observation is further supported by our finding that there was no difference in the average multivariate plastic response to temperature in our three populations, but there were differences in the population’s average multivariate plastic response to food. This finding appears to contradict previous studies that used laboratory experiments, mesocosm experiments and resurrection ecology to demonstrate that in *D. magna* populations, thermal tolerance is genetically variable and can evolve rapidly in response to increased temperatures (Van Doorslaer *et al*., 2007, 2009a, b, 2010; De Meester *et al*., 2011; Geerts *et al*., 2015). One explanation could be that in previous studies thermal tolerance evolved as an indirect consequence of selection for another trait (De Meester *et al*., 2011). Geerts *et. al*. (2017) demonstrated that CTmax was negatively genetically correlated with body size. Our study also found a negative genetic correlation between CTmax and the average number of offspring per clutch, a trait that is closely associated with body size (see Fig. 2, 3). So it is possible that selection for body size and/or faster demographic rates (Bruijning *et al*., 2018) explained the evolution of thermal tolerance in previous studies. This interpretation might also explain why temperature manipulations that are still in many cases at least 10°C below CTmax values are capable of generating rapid adaptation in experimental populations (van Doorslaer *et al*., 2009b). Our finding that increases in resource availability generate plastic decreases in CTmax (Fig. 4) but changes is temperature do not induce plastic shifts in CTmax (Fig. 3), also support the idea that thermal tolerance evolves indirectly. We therefore suggest that a multivariate understanding of rapid adaptation to thermal environments is required before we can determine whether the rapid evolution of thermal tolerance reported in numerous studies is a direct result of selection for thermal tolerance, or an indirect consequence of the effect that temperature has on the evolutionary potential of traits such as body size and population growth rate.

In summary, we have demonstrated that multivariate plastic responses to food and temperature explained three times more phenotypic variation than genetic variation in traits or trait plasticities. G x E interactions exist in natural populations of *Daphnia magna* but they are typically context-dependent. For temperature plasticity, the context-dependence manifests as a shift in the suite of traits that explain the most additive genetic variance in different food environments. But for food plasticity, the context-dependence also resulted in a reduction in evolutionary potential in low food at 18°C. This reduced evolutionary potential may explain why the population still harbours additive genetic variation in traits related to adaptive plastic responses to food, but little additive genetic variation in traits involved in adaptive plastic responses to temperature.

## ACKNOWLEDGEMENTS

We thank Luc De Meester for the provision of *Daphnia* clones from France and Denmark and Aurora Geerts for advice on the heat tolerance assay. The study was supported by NERC Highlight grant NE/N016017/1 to SJP, DA & SPa.

## DATA ACCESSIBILITY

Data will be archived in Dryad upon acceptance.

## Notes

### Competing Interest Statement

The authors have declared no competing interest.

