## Supplementary tables and figures for "Quantifying multivariate genotype-by-environment interactions, evolutionary potential and its context-dependence in natural populations of the water flea, *Daphnia magna*"

**Supplementary Methods**

In addition to the base R version 3.5.2 (R Core Team 2018), we used the packages car (Fox and Weisberg 2011), ggplot2 (Wickham 2016), Hmisc (Harrell Jr 2019a), lme4 (Bates et al. 2015), MASS (Venables and Ripley 2002), rms (Harrell Jr 2019b), SDMTools (VanDerWal et al. 2014), vegan (Oksanen et al. 2018), VennDiagram (Chen 2018) and R code provided in Collyer & Adams (2007).

To assess multivariate phenotype differences between treatments, as well as between population variation values, we analysed the 8 life history variables together with the thermal tolerance of each individual in a perMANOVA. To this end, pairwise Gower distances between multivariate phenotypes were first calculated with vegdist {vegan}. This distance matrix was then assessed in a perMANOVA with 9999 permutations using the adonis {vegan} function, setting temperature, food levels, population and all interactions as fixed factors, with temporal block described as a random factor. The same analysis was repeated within each population to test for clonal variation in responses, setting clone instead of population as a fixed factor.

To calculate components of variation in data subsets, e.g. for the temperature response data of the UK population in a low food context, perMANOVAs as described above were carried out on the relevant data subset, and rerun with different orders of the tested factors to obtain marginal R^2^ estimates for each tested factor.

We calculated magnitude and phenotypic integration plasticity values for the complete dataset as well as within environmental context, i.e. average temperature responses across data from both food treatments and temperature responses in a high and low food environment, using the scaling for the full data set from the respective population throughout. In the same way, life history plasticity in response to food levels was calculated across all data and separately for the high and low temperature environments.

To visualise multivariate phenotypes, we used principal component analyses (PCAs) on scaled data for each population, using the prcomp {stats} and ordiplot {vegan} functions. Theoretical plasticity distributions in multidimensional traits space (Fig. 1) were plotted by changing the standard variation around the same mean reaction norm for either the reaction norm length, the reaction norm angle or the reaction norm midpoint, when randomly drawing those values from a normal distribution

**Table S1: perMANOVA results across populations**

| **factor** | **Df** | **SofSqu** | **MeanSqus** | **F-value** | **R2** | **P** |
| --- | --- | --- | --- | --- | --- | --- |
| **T** | 1 | 4.327 | 4.327 | 440.774 | 0.234 | **<0.001** |
| **food** | 1 | 5.327 | 5.327 | 542.661 | 0.288 | **<0.001** |
| **Pop** | 2 | 0.415 | 0.208 | 21.162 | 0.022 | **<0.001** |
| **T:food** | 1 | 0.091 | 0.091 | 9.223 | 0.005 | **<0.001** |
| T:Pop | 2 | -0.030 | -0.015 | -1.545 | -0.002 | 1.000 |
| **food:Pop** | 2 | 0.047 | 0.023 | 2.381 | 0.003 | **0.017** |
| T:food:Pop | 2 | 0.020 | 0.010 | 1.019 | 0.001 | 0.344 |
| Residuals | 844 | 8.285 | 0.010 |  | 0.448 |  |

significant results are highlighted in bold

**Table S2: perMANOVA results within populations**

| **a) UK** | **Df** | **SofSqu** | **MeanSqus** | **F-value** | **R2** | **P** |
| --- | --- | --- | --- | --- | --- | --- |
| **T** | 1 | 5.048 | 5.048 | 639.012 | 0.286 | **<0.001** |
| **food** | 1 | 4.755 | 4.755 | 601.955 | 0.269 | **<0.001** |
| **Clone** | 42 | 2.115 | 0.050 | 6.376 | 0.120 | **<0.001** |
| T:food | 1 | -0.013 | -0.013 | -1.688 | -0.001 | 1.000 |
| **T:Clone** | 42 | 0.828 | 0.020 | 2.496 | 0.047 | **<0.001** |
| **food:Clone** | 42 | 0.544 | 0.013 | 1.639 | 0.031 | **<0.001** |
| **T:food:Clone** | 42 | 0.446 | 0.011 | 1.345 | 0.025 | **0.015** |
| Residuals | 501 | 3.958 | 0.008 |  | 0.224 |  |

| **b) F** | **Df** | **SofSqu** | **MeanSqus** | **F-value** | **R2** | **P** |
| --- | --- | --- | --- | --- | --- | --- |
| **T** | 1 | 1.052 | 1.052 | 85.581 | 0.327 | **<0.001** |
| **food** | 1 | 0.933 | 0.933 | 75.861 | 0.289 | **<0.001** |
| **Clone** | 4 | 0.209 | 0.052 | 4.250 | 0.065 | **<0.001** |
| T:food | 1 | 0.018 | 0.018 | 1.490 | 0.006 | 0.211 |
| T:Clone | 4 | 0.061 | 0.015 | 1.244 | 0.019 | 0.284 |
| **food:Clone** | 4 | 0.117 | 0.029 | 2.382 | 0.036 | **0.009** |
| **T:food:Clone** | 4 | 0.119 | 0.030 | 2.413 | 0.037 | **0.010** |
| Residuals | 58 | 0.713 | 0.012 |  | 0.221 |  |

| **c) DK** | **Df** | **SofSqu** | **MeanSqus** | **F-value** | **R2** | **P** |
| --- | --- | --- | --- | --- | --- | --- |
| **T** | 1 | 0.653 | 0.653 | 93.273 | 0.213 | **<0.001** |
| **food** | 1 | 1.051 | 1.051 | 150.200 | 0.342 | **<0.001** |
| **Clone** | 7 | 0.462 | 0.066 | 9.430 | 0.150 | **<0.001** |
| **T:food** | 1 | 0.083 | 0.083 | 11.904 | 0.027 | **<0.001** |
| T:Clone | 7 | 0.077 | 0.011 | 1.581 | 0.025 | 0.082 |
| **food:Clone** | 7 | 0.194 | 0.028 | 3.966 | 0.063 | **<0.001** |
| T:food:Clone | 7 | 0.038 | 0.005 | 0.768 | 0.012 | 0.731 |
| Residuals | 73 | 0.511 | 0.007 |  | 0.166 |  |

significant results are highlighted in bold

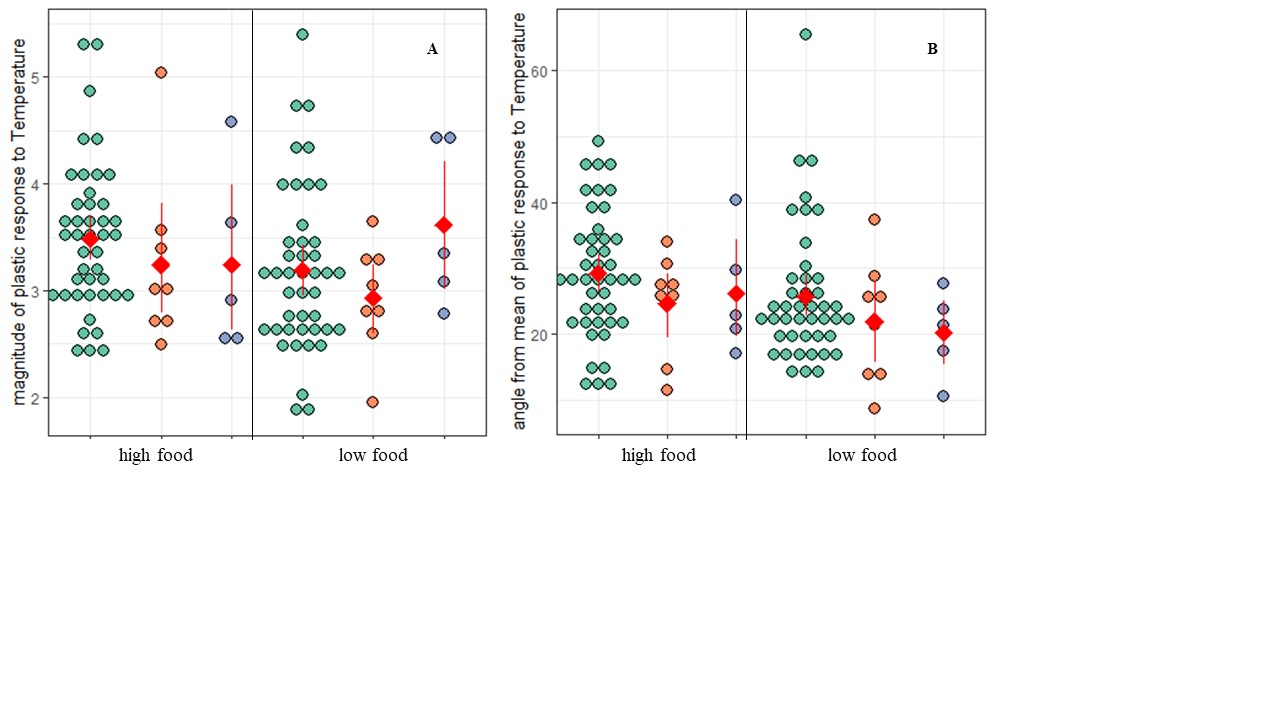

**Figure S1: Variation in magnitude and phenotypic integration of plastic responses in 3 different populations.**

Plastic responses of individual clones to temperature difference in separate food environments, measured as Euclidian distance of reaction norm in multivariate trait space (A) and phenotypic integration differences from the respective population mean for individual clones, measured as angle between reaction norm of individual clone and the average reaction norm of the respective population (B). Clone values are grouped by population: green = UK, orange = DK, purple = F. Population means and non-parametric bootstrap confidence intervals are shown in red.
